# New climatically specialized lineages of *Batrachochytrium dendrobatidis* and their sub-lethal effects on amphibians establish the Asiatic origins of the pathogen

**DOI:** 10.1101/2023.01.23.525302

**Authors:** Dan Sun, Gajaba Ellepola, Jayampathi Herath, Hong Liu, Yewei Liu, Kris Murray, Madhava Meegaskumbura

**Affiliations:** Guangxi Key Laboratory for Forest Ecology and Conservation, College of Forestry, Guangxi University; Nanning, Guangxi 530000, People’s Republic of China; Department of Zoology, Faculty of Science, University of Peradeniya, Peradeniya, KY20400, Sri Lanka; Guangxi Huaping Natural Nature Reserve Administration, Guangxi 530000, People’s Republic of China; MRC Unit The Gambia at London School of Hygiene and Tropical Medicine, Atlantic Boulevard, Fajara, The Gambia

**Keywords:** Fungal disease, Chytridiomycosis, *Batrachochytrium dendrobatidi*, Evolution, Origin, Environmental conditions

## Abstract

Chytridiomycosis, a highly significant global wildlife disease, has caused unprecedented amphibian population declines and species extinctions worldwide. In contrast, mass die-offs due to chytridiomycosis have not been observed in Asia, which is thought to be the ancestral region of origin and a hyper-diversity hotspot of the known causal pathogens, *Batrachochytrium dendrobtidis* (*Bd*) and *B. salamndrivorans* (*Bsal*). It has been hypothesized that Asian amphibians may have evolved immunity to clinical *Batrachochytrium* infection. However, limited knowledge of endemic lineages, evolutionary history, and climate-related infection patterns limits our ability to explore this hypothesis. Here, we investigated the genetic diversity and infection patterns of the frog-infecting species, *Bd*, in China’s poorly-explored Guangxi region. We used the internal transcribed spacer (ITS) marker and the nested PCR method to survey prevalence and haplotype diversity of *Bd* across 17 forest sites. A generalized linear model was used to evaluate associations between numerous variables and *Bd* prevalence within native amphibians. Our results identified seven new haplotypes, four of which are closely related to the early-emerging *Bd*ASIA-1 lineage recovered from South Korea. We also identified a unique Asian haplotype, close to the *Bd*ASIA-3 lineage, as the most prevalent (64.6% of *Bd*-infected adult individuals) in 11 out of 15 infected species. This haplotype was also detected in a salamander individual, which exhibited non-lethal skin lesions on the abdomen. The infection of *Bd* within amphibians was found to be positively associated with temperature and elevation. Our findings suggest that there is significant undiscovered genetic diversity of Asian *Bd* lineages in this region. Longer-term studies are required to further investigate *Bd* diversity, prevalence, seasonality and impact on native species and populations in Southern China and across the region of origin in Asia.

**Author Summary:** Chytridiomycosis is a disease which is responsible for the sharp decline of amphibian populations and species extinctions around the world. Surprisingly, it has not yet been well-studied in Asia, the region where the two causal pathogens of the disease, *Batrachochytrium dendrobatidis* (*Bd*) and *B. salamandrivorans* (*Bsal*) originated. In order to better understand the lack of mass die-offs in Asia, we recently conducted a study in south China’s Guangxi region to investigate the genetic diversity and infection patterns of *Bd*. Through the use of internal transcribed spacer (ITS) markers and nested PCR, we discovered seven new types of *Bd*, four of which were closely related to the early-emerging *Bd*ASIA-1 lineage from South Korea. The highest prevalence of *Bd* infection was observed in 11 species of amphibians, including a salamander which had non-lethal skin lesions. It was also noted that infection of *Bd* in amphibians was associated with temperature and elevation. This study has provided important information on *Bd* diversity and prevalence in the region, and further research is needed to explore Asia as the putative region of origin for this disease.

## Introduction

Emerging pathogenic fungi, such as *Batrachochytrium dendrobatidis* (*Bd*) and *B. salamndrivorans* (*Bsal*), pose a significant threat to many free-ranging animal taxa, particularly amphibians [1–3]. Since their discovery in the late 1990s and 2013, respectively, these highly virulent pathogens have been known to cause chytridiomycosis in amphibians, resulting in unprecedented impacts on global amphibian diversity [4–9]. In particular, *Bd* is considered one of the most feared fungal pathogens infecting vertebrate taxa [8, 10]. It is thought that the geographic origin of *Bd* is in Asia [11]; however, mass mortality of amphibians has not been recorded in this region [12, 13]. Despite this, our understanding of the endemic lineages, complex evolutionary history, genetic diversity, infection patterns, and predictors of *Bd* in the region of origin remains limited.

Asia is considered a “cold spot” for chytridiomycosis, as the presence of *Bd* has not been associated with mass die-offs or amphibian population declines [13, 14]. Moreover, prevalence (below 10%) and infection loads (zoospore loads <10000) in infected Asian amphibians are usually below the thresholds associated with clinical disease in other regions [15–17]. It is possible that native amphibian species in Asia have evolved resistance to the clinical impacts of *Bd* infection over the past 50 million years, due to the presence of ancient *Bd* in natural urodele hosts, such as *Andrias japonicas* [6, 7, 18, 19].

Nevertheless, some Asian amphibian populations have been found to exhibit higher prevalence of *Bd* (i.g. >20%) [18, 20–22]. These populations often harbor multiple *Bd* genotypes, including both endemic lineages and the globally invasive *Bd*GPL. This suggests that certain species in the region may serve as important reservoir hosts, allowing the pathogen to persist across the region [18, 21–23].

Two endemic *Bd* lineages, *Bd*ASIA-1 and *Bd*ASIA-2/*Bd*BRAZIL, have been identified in South Korea [11, 22], and *Bd*ASIA-3 has recently been described from Indonesia, Philippines, and China [23]. High ITS-based haplotype diversity of *Bd*GPL and four specific early-emerging haplotypes (JPK, JPJ, IN55, and JPB=CN30=IN05) have been identified in Japan, India, and the Yunnan province of China [18, 23, 24], suggesting that a wide area within the Asian continent is likely the area of origin of *Bd* and the main driver of its genetic diversity.

Although China has the largest number of amphibian species in Asia (more than 500 species) [25], only 110 species have been tested for *Bd* infection [26–28]. Of these, approximately 30 native species, as well as two additional exotic species (African clawed frog *Xenopus laevis* and the American bullfrog *Rana catesbeiana*), have been confirmed positive. Despite this, understanding of the diversity and evolution of native *Bd* lineages and their host dynamics is poor in China, likely due to the narrow distributional ranges of many native species [25]. Infected native amphibians have mostly been recorded from the southern regions of China [24, 26–30], which is characterized by a complex geography and pronounced climate variation, particularly in natural forested mountains. Therefore, further knowledge of genetic diversity and the environmental correlates of *Bd* distribution and prevalence in affected habitats across China is essential [23, 24, 30, 31].

We hypothesized (1), that southern China likely harbours undiscovered genetic diversity of both endemic Asian *Bd* lineages and globally invasive lineages infecting previously undocumented host species, and (2), that environmental factors, particularly habitat and climate, play a key role in structuring this diversity and infection prevalence. To explore this, we conducted a widespread genetic sampling of *Bd* lineages across 17 sites consisting of various environments and native species.

We found new basal haplotypes that putatively belong to the *Bd*ASIA-1 and *Bd*ASIA-3 lineages, suggesting that China is an important epicenter of evolutionary process for *Bd* and highlighting the need for further in-depth studies of *Bd* diversity and infection patterns in the region.

## Materials and methods

### Study sites

The Guangxi Autonomous Region (GAR, 21°42.45’-25°37.01’ N, 107°32.59’-110°12.44’ E), located in southern China (Fig. 1), covers 236,700 km^2^ and is home to a mosaic of habitats that host amphibian species. *Bd* has been reported in seven amphibian species from GAR: a wild caecilian (*Ichthyophis bannanicus*) [28], five frog species (*Fejervarya limnocharis, Odorrana schmackeri, Rhacophorus dennysi, Rhacophorus megacephalus, Amolops ricketti*) [30, 33], and one toad species (*Bufo melanostictus*) [29, 32]. The western border of GAR is shared with the province of Yunnan, from which an endemic Asian *Bd* lineage has been described [23].

**Fig 1.**
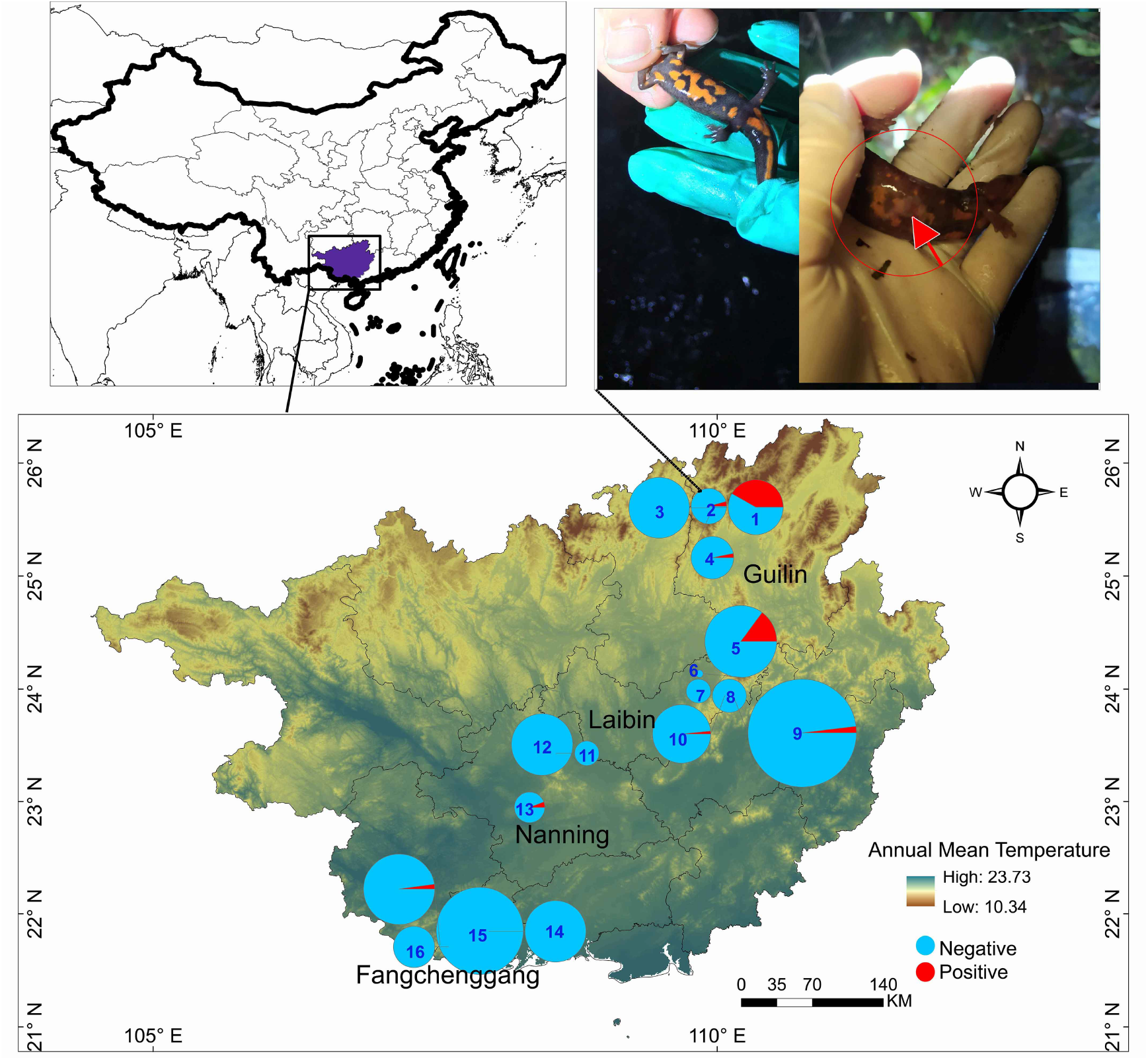
The distribution of sampling sites across a latitudinal gradient. The map depicts annual mean temperature of the study region. Fangchenggang, Nanning, Laibin and Guilin represent four cities where sampling sites located. Numbers represent different sampling sites: 1 (Hongtan), 2 (Huaping), 3 (Cujiang), 4 (Anjiangping), 5 (Shiliugongli), 6 (Hekou), 7 (Laoshanshuiku), 8 (Wuzhishan), 9 (Shengtangshan), 10 (Daling), 11 (Shangshuiyuan),12 (Tianping), 13 (Laohuling), 14 (Nakuan), 15 (Pinglong), 16 (Nianbei), 17 (Dongzhong). The size variations of the circles correspond to the numbers of skin swab samples from adults; – further details are provided in the supplemental materials (S1 Table). An abdomen of a healthy *P*. *inexpectatus* salamander (left) and observed skin lesions on the abdomen of a *P. inexpectatus* salamander (right), infected by *Bd* at Site #2. The location of the study region, Guangxi in southern China.

We focused our sampling on four natural and protected forest areas along a latitudinal gradient across GAR (S1Table). These areas lie within the Indo-Burma biodiversity hotspot [33] and capture the climatic and topographic variation of Southern China. The region’s climate is sub-tropical and depends on the central and South Asian tropical monsoon, with strongly seasonal rainfall [34]. Our sampling sites ranged in altitude ca. 70 – 1300 m, with vegetation dominated by sub-tropical evergreen broad-leaved forests in the north and sub-tropical evergreen seasonal rainforests in the south.

Seventeen field sites were selected based on information on amphibian habitats and accessibility. Seven of these were in southern Guangxi and the remaining ten were in northern Guangxi (Fig. 1). The sites spanned an altitudinal range from 69 – 1310 m above sea level (a.s.l.) and had varying levels of disturbance due to human activities such as villages, tourism, and agriculture. Most of the sites were separated by more than 5 km, but 6 sites were less than 5 km apart because of impassable geographical barriers, such as mountains and torrents.

We employed nocturnal visual encounter and acoustic methods to collect *Bd* samples from amphibians between 2019 and 2021. Samples were collected from amphibians dwelling in various habitat types, including forest floors, tree branches, leaves and trunks, and various water bodies such as permanent lakes, periodic and perennial ponds, streams, riparian zones, and ephemeral habitats. We also conducted opportunistic sampling of tadpoles when possible. The infection status of wild amphibians was evaluated by assessing typical clinical signs, including skin lesions, coloration, and shedding of adult amphibians [4, 35, 36], and by looking for loss or deformities in keratin-covered regions such as mouthparts, tooth rows, and jaw sheaths for larval anurans [37, 38].

### Molecular diagnostics of *Bd*

We swabbed 1088 amphibians consisting of 1012 adults and 76 tadpoles. They consisted of 36 species across 25 genera (S1 Table). All sampling adhered to approved protocols from the Institutional Animal Care and Use Committee (GXU2018-048) at Guangxi University and was conducted with ethical clearance.

We collected individuals using clean and unused 10 × 5 cm or 15 × 20 cm plastic zippered bags. We followed standardized protocols and biosecurity measures while collecting skin and mouthpart swabs from wild amphibians. To collect mouthpart swabs from larval amphibians, we gently inserted a sterile dry swab with a fine-tip (Medical Wire & Equipment Co. MW 113) and rotated it fifteen times. For adult amphibians, we rubbed their skin with a swab using some pressure and firmly moving the swab over the back, pelvic patch, inside back legs and toes ten times [39–42]. After swabbing, we immediately sealed the swabs into 1.5 ml Eppendorf tubes without touching and released the amphibians back to their point of capture. We stored the swab samples at −80°C in the lab until DNA extraction.

DNA extraction from swab samples were carried out using PrepMan Ultra reagent (Applied Biosystems, Foster City, CA) [43] and Qiagen DNeasy Blood and Tissue Kit [44]. The extracted genome DNA was diluted with nuclease-free water in proportions of 1:10 for follow-up molecular diagnosis, and stored at −80°C [41, 45].

We used nested PCR assay for the detection of *Bd* [18, 46]. Total 25μl reactions contained 12.5μl, 0.5nM of each primer, 5μl DNA template. The conditions for the first amplification included initial denaturation for 4min at 94°C, 30 cycles of 30s at 94°C, 30s at 50°C and 2min at 72°C, and a final extension for 10min at 70°C. The conditions for the second amplification consisted of initial denaturation for 4min at 94°C, 30 cycles of 30s at 94°C, 30s at 60°C and 1min at 72°C and a final extension for 10min at 72°C. PCR products were visualized on 1.5% agarose gels, around 300 bp bands were observed, and positive samples for *Bd* were confirmed via re-amplification. Negative and positive controls were used in each PCR amplification. PCR products of positive samples from the second amplification were directly sequenced though Sequencer (Sangon Biotech). A few of the sequenced samples with ambiguous chromatograms of amplicon and positive samples with multiple bands were cloned using Hieff Clone® Zero TOPO-TA Cloning Kit in accordance with the manufacturer’s protocol and sequenced using universal primers.

### Data analysis

#### Variations in *Bd* prevalence

We initially summarized infection patterns across *Bd* prevalence and corresponding 95% confidence intervals according to age (adults and larvae), species, sites and months. We used the ‘binconf’ function with Wilson interval in the “Hmisc” package [47]. Then, we further analyzed positive samples phylogenetically and assessed patterns of infection prevalence using generalised linear models (GMLs). See separate sections below for details.

#### Drivers of infection prevalence analysis

We employed an information-theoretic modelling approach [48] to assess the effects of multiple bioclimatic, elevation, season (month), habitat factors, and latitude □ longitude interaction on *Bd* infection in amphibian adults [49–54]. We downloaded 19 bioclimatic variables from the CHELSA version 1.2 database at a resolution of 30 arc sec [55]. We calculated the correlation between bioclimatic factors and elevation, and only selected five variables with a correlation coefficient < 0.70. To classify adult habitats, we used the activity breadth of adults observed during the non-breeding season [56, 57].

We conducted a Generalised Linear Model (GLM) to analyze the influence of eight predictor variables on *Bd* presence/absence in amphibian populations. We used populations infected by *Bd* within sites as the response variable. We set up candidate models based on all possible combinations of the eight factors as explanatory variables, as well as a null model. We also included a candidate model with species as the single explanatory variable to assess whether species per se affect *Bd* presence/absence.

Each candidate model was quantified and evaluated based on Akaike Information Criterion (AIC), Akaike second-order corrected (AICc) and Akaike weights (AICw) [58]. The final support model was validated according to the evaluation of homogeneity in the residuals of the models against fitted values [59].

#### Phylogenetic analysis

We analyzed the phylogenetic relationships among newly generated *Bd* haplotypes from this study combined with sequences of ITS1-5.8S-ITS2 gene generated in previous studies (S2 Table). Seven outgroup taxa were selected according to [18]. We aligned a total length of 300 bp sequences of ITS1-5.8S rRNA-ITS2 of *Bd* using Clustal W with manual adjustment in MEGA v.7 [60]. Bayesian phylogenetic inference was performed in MrBayes v.3.2.6 [61] with an evolutionary HKY+G model generated in jModelTest v.2.1.10 using the Bayesian Information Criteria (BIC) [62]. Bayesian analyses were run for 15 million generations in four MCMC chains and 1000 generations sampled, with the initial 25% discarded as burn-in. Running results were visualized to check effective samples sizes (ESS) in Tracer v.1.6 [63]. The trees and posterior probabilities for label nodes were visualized in FigTree v.1.4.3 [64]. Haplotype sequence data of ITS-5.8S gene were generated by calculating the number and diversity of haplotypes in DnaSP v.6 [65]. Haplotype networks were obtained from Network v.10 using the median-joining method [66].

Furthermore, we used the bioclimatic and elevation parameters to examine the environmental conditions among *Bd* haplotypes based on a principal component analysis method.

## Results

### Variation in prevalence of *Bd*

The overall prevalence of *Bd* infection across the GAR was 4.74% (95%CI: 3.60%-6.23%) in adult individuals and 5.26% (95%CI: 2.07%-12.77%) in tadpoles. Of 36 amphibian species sampled, 16 had positive *Bd* infections (Table 1). The highest prevalence among species with at least 22 samples was 59.09% (95%CI: 38.73%-76.74%) for *Amolops chunganensis*. Among a sample of 15 tadpoles from five or six species, 4 from *Hylarana guentheri* were *Bd* positive (26.67%, 95%CI: 10.90%-51.95%; Table 1).

**Table 1.** Prevalence of *Bd* infection and haplotype distribution in 16 infected species across 13 genera, with *Bd* haplotypes and numbers of haplotypes, and 95% CI are indicated within brackets. ✶ indicate *Bd* positive species from previous studies in Asia. Conservation status is based on IUCN categories with acronyms representing from low to high extinction risk: LC, least concern; NT, near threatened; VU, vulnerable; EN, endangered; DD, Data Deficiency.

| Genus | Species | Sample | Positive (Haplotype) | Prevalence (95%CI) | IUCN |
| --- | --- | --- | --- | --- | --- |
| *Amolops* | Chungan Sucker Frog, *Amolops chunganensis* | 22 | 13 (GX03) | 59.09% (38.73%-76.74%) | LC |
|  | South China Torrent Frog, *Amolops ricketti* | 271 | 1 (GX10) | 0.37% (0.07%-2.06%) | LC |
| *Kurixalus* | Serrate-legged Small Treefrog, *Kurixalus odontotarsus* | 42 | 2 (1: GX02; 1: GX03) | 4.76% (1.32%-15.79%) | LC |
| *Odorrana* | Lungshen Odorous Frog, *Odorrana lungshengensis* | 22 | 6 (GX03) | 27.27% (13.15%-48.15%) | NT |
|  | Large Odorous Frog, *Odorrana graminea* | 53 | 2 (GX03) | 3.77% (1.04%-12.75%) | DD |
|  | Bamboo-leaf Frog, *Odorrana versabilis* | 20 | 3 (GX03) | 15.00% (5.24%-36.04%) | LC |
| *Pachytriton* | Yaoshan Stout Newt, *Pachytriton inexpectatus* | 20 | 2 (GX03) | 10.00% (2.79%-30.10%) | LC |
| *Quasipaa* | Spiny-bellied Frog, *Quasipaa boulengeri*★ | 34 | 1 (GX03) | 2.94% (0.52%-14.92%) | EN |
| *Rana* | Hanlui Brown Frog, *Rana hanluica* | 4 | 1 (GX03) | 25.00% (4.56%-69.94%) | DD |
| *Kaloula* | Asian Painted Frog, *Kaloula pulchra*★ | 1 | 1 (GX04) | 100% (20.65%-100.00%) | LC |
| *Polypedates* | Spot-legged Treefrog, *Polypedates megacephalus*★ | 131 | 1 (GX03) | 0.76% (0.13%-4.20%) | LC |
| *Zhangixalus* | Large Treefrog, *Zhangixalus dennysi*★ | 32 | 1 (GX03) | 3.13% (0.55%-15.74%) | LC |
| *Leptobrachella* | Fujian Metacarpal-tubercled Toad, *Leptobrachella liui* | 19 | 3 (1: GX01; 2: GX05) | 15.79% (5.52%-37.57%) | LC |
| *Theloderma* | Red-disked Small Treefrog, *Theloderma rhododiscus* | 38 | 8 (7: GX05; 1: GX03; 1: GX06) | 21.05% (11.07%-36.35%) | NT |
| *Zhangixalus* | Minimal Treefrog, *Zhangixalus minimus* | 37 | 3 (2: GX05; 1: GX09) | 8.11% (2.80%-21.30%) | EN |
| *Hylarana* | Guenther's Stream Frog<br>*Hylarana guentheri* (Larva) | 15 | 4 (1: GX07; 1: GX08; 2: GX01) | 26.67% (10.90%-51.95%) | LC |

Dead or dying frogs affected with *Bd* were not observed in the field. However, the typical clinical signs of chytridiomycosis were noted in one individual of a Yaoshan Stout Newt (*Pachytriton inexpectatus*), which contained abdominal skin lesions (Fig. 1).

The prevalence of *Bd* varied between sites (range 0.0% to 43.4%; see Table 2 and Fig. 1). Sites #1 (Hongtan) and #5 (Shiliugongli), both located at high elevations (>700 m), had the highest prevalence. Sample sizes and infection status are indicated in S1 Table. *Bd* prevalence was higher in northern GAR sites during the colder, rainy season (S3 Table).

**Table 2.** Prevalence of *Bd* and haplotype distribution in the 17 sampling sites. *Bd* haplotypes with their numbers, and 95% CI are indicated within brackets.

| Site # (location: elevation) | No. sample | No. positive (haplotype) | Prevalence (95%CI) |
| --- | --- | --- | --- |
| Site#1 (Hongtan: 889 m) | Adult: 53 | 23 (GX03) | **43.40% (31.95%-56.73%)** |
| Site #2 (Huaping: 1033 m) | Adult: 30 | 1 (GX03) | 3.33% (0.59%-16.67%) |
| Site #3 (Cujiang: 965 m) | Adult: 54<br>Larva: 15 | 4 (2: GX01; 1: GX07; 1: GX08) | 5.80% (2.27%-13.98%) |
| Site #4 (Anjiangping: 1310 m) | Adult: 43<br>Larva: 16 | 1 (GX03) | 1.69% (0.30%-8.90%) |
| Site #5 (Shiliugongli: 1286 m) | Adult: 102<br>Larva: 8 | 14 (1: GX01; 1: GX03; 11: GX05; 1: GX06; 1: GX09) | **12.72% (7.73%-20.24%)** |
| Site #6 (Hekou: 841 m) | Adult: 1<br>Larva: 29 | 0 | 0.00% (0.00-11.35%) |
| Site #7 (Laoshanshuiku: 943 m) | Adult: 11 | 0 | 0.00% (0.00%-25.88%) |
| Site #8 (Wuzhishan: 573 m) | Adult: 21 | 0 | 2.94% (0.52%-14.92%) |
| Site #9<br>(Shengtangshan: 770 m) | Adult: 214 | 4 (GX03) | 0.87% (0.73%-4.71%) |
| Site #10 (Daling: 476 m) | Adult: 62 | 1 (GX10) | 1.61% (0.29%-8.59%) |
| Site #11 (Shangshuiyuan: 69 m) | Adult: 11 | 0 | 0.00% (0.00%-25.88%) |
| Site #12 (Tianping: 1249 m) | Adult: 69 | 0 | 0.00% (0.00%-5.27%) |
| Site #13 (Laohuling: 104 m) | Adult: 17 | 1 (1: GX04) | 5.88% (1.05%-26.98%) |
| Site #14 (Nakuan: 131 m) | Adult: 68 | 0 | 0.00% (0.00%-5.35%) |
| Site #15 (Pinglong: 509 m) | Adult: 142 | 1 (GX03) | 0.70% (0.12%-3.88%) |
| Site #16 (Nianbei: 282 m) | Adult: 31<br>Larva: 2 | 0 | 0.00% (0.00%-10.43%) |
| Site #17 (Dongzhong: 912 m) | Adult: 92<br>Larva: 6 | 2 (1: GX02; 1: GX03) | 2.04% (0.56%-7.14%) |

### Drivers of *Bd* prevalence

The best-supported GLM involved two variables: mean temperature of the warmest quarter and elevation (S4 Table). This suggests that high temperature and high-low elevation were positively correlated with *Bd* prevalence within amphibians; other environmental and geographical factors had a marginal effect on *Bd* infection, but no effect was linked to species.

### Phylogenetic diversity

#### ITS-based haplotype diversity and distribution

A total of 10 haplotypes were detected in the 52 positive samples. Each infected amphibian had only a single haplotype, except for the tree frog species *Theloderma rhododiscus*, which harbored two haplotypes (GX05 and GX06). These haplotypes were numbered from GX01 to GX10 and deposited in GenBank under the accession numbers OQ275246 – OQ275255.

Seven of the ten *Bd* haplotypes were novel; the remaining three had been previously reported (S2 Table). Haplotype GX03 was identical to CN30 from Yunnan Province, China [24], JPB from Japan [18], and IN05 from mountains in India [67]. It was also the most common haplotype at all sites where *Bd* was found in this study. GX03 was detected in 32 adults from 11 species, representing 73% of the 16 susceptible species and 67% of the 48 infected adults (Table 1).

#### Phylogenetic relationships and haplotype network

Our phylogenetic analysis on *Bd* based on 254 ITS sequences revealed five major clades in the Bayesian phylogenetic tree (Fig. 2). Haplotypes GX04, GX05, GX06 and GX10 formed an independent cluster nested in the most basal clade, closely related to Korean haplotypes. GX03, GX07, GX08 and GX09, along with several previously identified haplotypes from Japan, China and India, formed the second major clade. The third and fourth clades included Brazilian, Japanese, South African and other haplotypes. The fifth clade was distinctive from the *Bd*GPL clade and contained Asian haplotypes, while GX01 and GX02 were placed in the fourth clade together with *Bd*GPL haplotypes.

**Fig 2.**
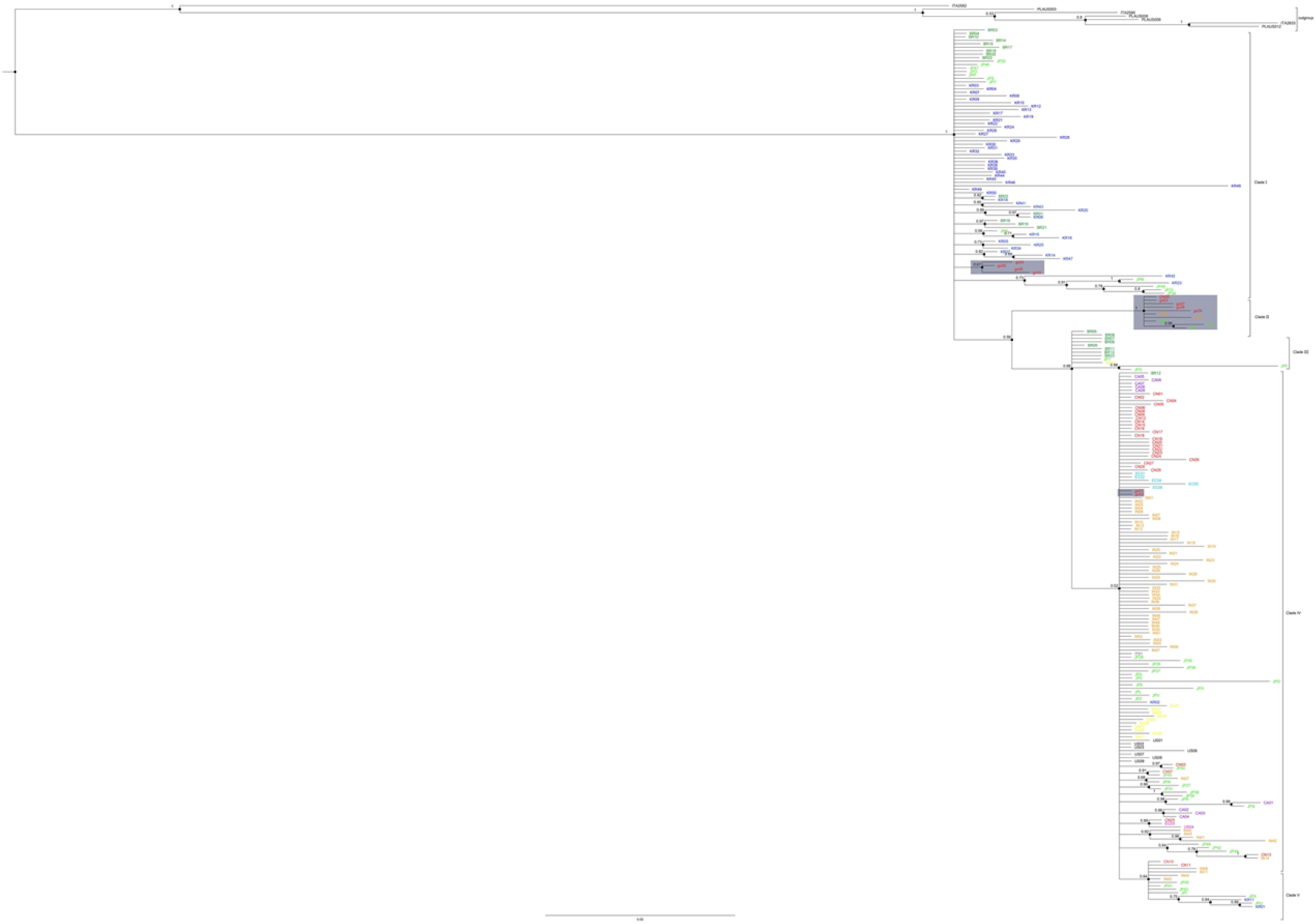
Phylogenetic tree of *Bd* ITS1-5.8S rRNA-ITS2 haplotypes based on Bayesian inference. The posterior probability (PP > 0.5) values are shown at nodes. Haplotypes marked with different colors indicates ten countries.

The median-joining network revealed 207 *Bd* haplotypes (10 newly generated and 197 obtained from GenBank) distributed geographically (Fig. 3 and S2 Table). The ten haplotype clusters found in this analysis were similar to the clades on the phylogenetic tree. Haplotypes GX04-GX06 and GX10 formed a separate cluster, likely belonging to the *Bd*ASIA-1 lineage recently identified in the fire-bellied toad *Bombina orientalis* from South Korea [11]. Interestingly, haplotypes GX03, GX07-GX09, combined with IN55, JPK and JPJ, formed another cluster which was distinct from the endemic *Bd* lineages, and was likely part of the *Bd*ASIA-3 lineage recently identified in Southeast Asia [23]. Haplotypes GX01 and GX02 were found within the group of the globally widespread *Bd*GPL lineage. Haplotype A (GX01) was the most prevalent haplotype across continents, while haplotype B (GX02) was detected in Ecuador, South Africa, Japan and China.

**Fig 3.**
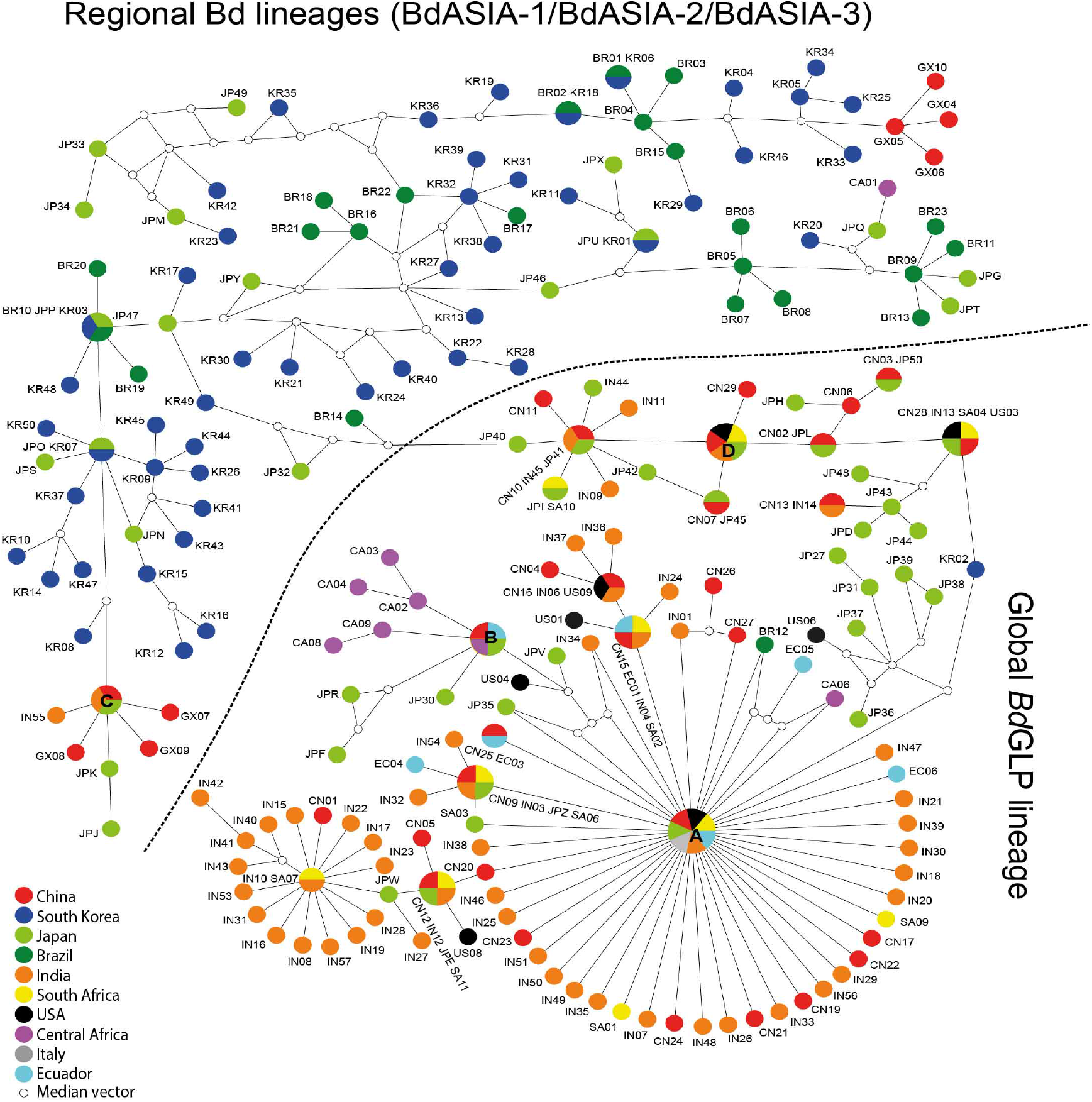
Median-joining haplotype network of *Bd* ITS1-5.8S rRNA-ITS2 sequences. Colored circles represent haplotypes from different countries. Hollow circles represent median vectors (missing haplotype). Owing to space limitations, the figure does not show the specific numbers of mutations in each step and denote same haplotypes instead of these abbreviations: A = CA05 = CN18 = IN02 = IT01 = JPA = SA08 = US02 = GX01; B=CA07 CN14 = EC02 = JPC = GX02; C = CN30 = IN05 = JPB = GX03; D = CN08 = IN52 = JP29 = SA05 = US07.

#### Putative environmental conditions of *Bd* lineages found in the study

Principal components analysis of bioclimatic and elevation parameters for *Bd* lineages (Fig. 4) revealed that the first two axes explained 93% of contribution. The *Bd*GPL and *Bd*ASIA-3 lineages were concentrated in the first axis, with the highest contributions from mean diurnal range, temperature seasonality, and precipitation of the warmest quarter. *Bd*ASIA-1, however, showed clustering in the second axis, with the highest contributions from mean temperature of the warmest quarter and elevation. These results suggest that different lineages occur under distinct climatic conditions.

**Fig 4.**
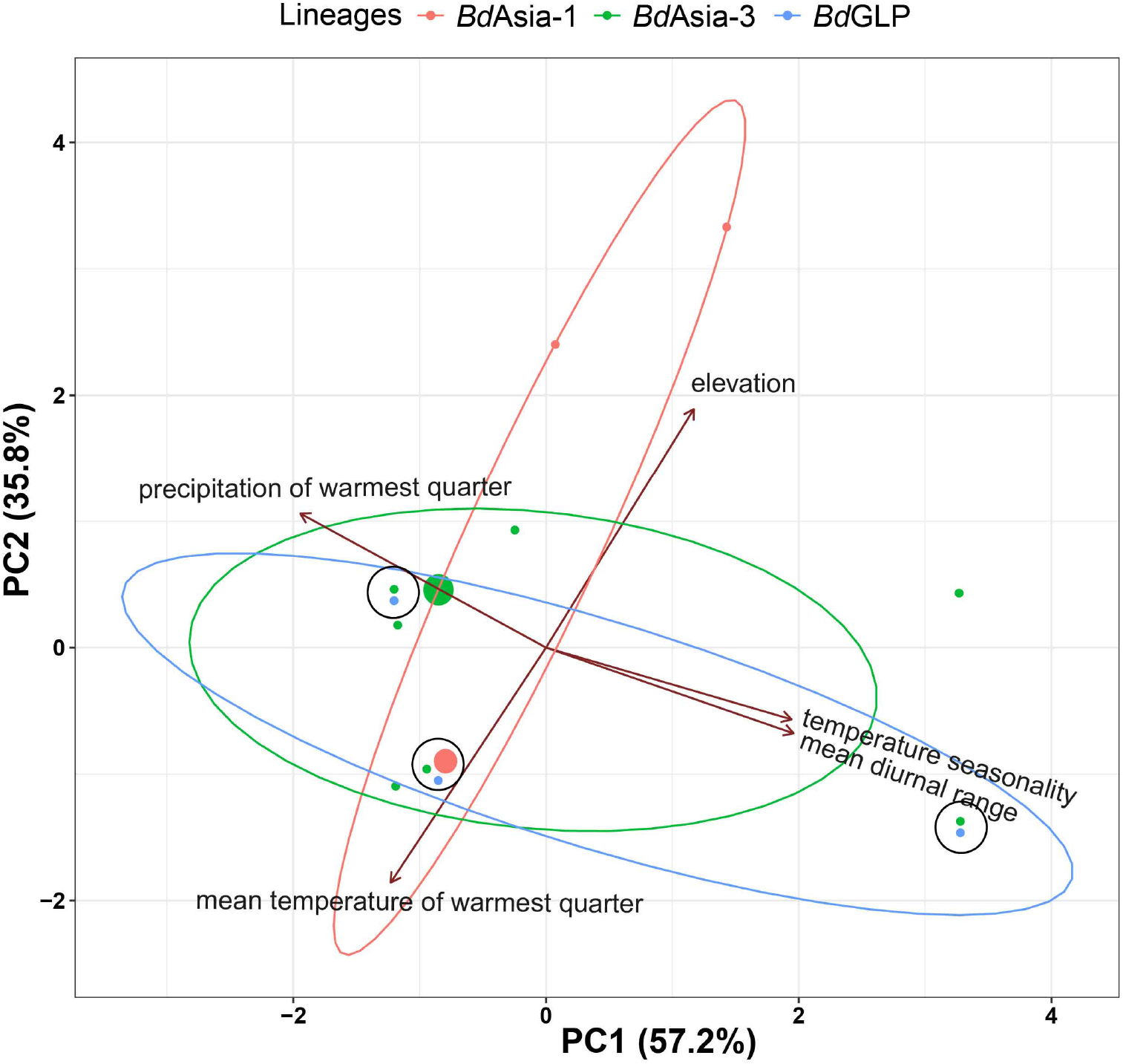
Principal components analysis of bioclimatic and elevation variables for *Bd* lineages. Different color ellipses represent each lineage. Black circles represent sites where two or three lineages co-exist. Points with different color represent sites where lineages exist, in which bigger green and red points represent high prevalence for *Bd*ASIA-3 and *Bd*ASIA-1 lineages, respectively.

## Discussion

Our hypothesis that there is undiscovered diversity of *Bd* lineages across South China was supported by our explorations across GAR. We discovered four new *Bd* lineages close to *Bd*ASIA-1, infecting three species in isolated populations. This indicates that the *Bd*ASIA-1 lineage has a broader distribution in Asia than previously thought. In addition, we identified four *Bd* haplotypes closely related to *Bd*ASIA-3, which dominate montane forests in southern China. Our findings suggest that the habitats of these *Bd* lineages are correlated with climatic and elevation factors, possibly explaining their localized distributions.

### Patterns of *Bd* infection

We identified 12 new amphibian species infected with *Bd*, in addition to the four species previously documented from different areas. This indicates that a greater number of forest amphibians in China are infected with *Bd* than was previously thought. The four previously recorded species (*K. pulchra, P. megacephalus, Q. boulengeri, Z. dennysi*) are all endemic to South-East Asia [25, 29, 32, 68].The newly identified hosts mainly occupy restricted regions of South-East Asia or China [25]. Our study screened 36 species from 25 genera, which is approximately one-third of all amphibian species from GAR. Notably, with the increasing number of newly described amphibian species from China (see, http://www.amphibiachina.org/), it is likely that numerous other restricted species are infected with *Bd*.

Our results suggest that some amphibian species and populations are highly susceptible to *Bd* infection. The Chungan Sucker Frog (*A. chunganensis*) exhibited the highest prevalence, while the Lungshen Odorous Frog (*O. lungshengensis*) and Red-disked Small Treefrog (*T. rhododiscus*) also showed relatively high prevalence. These findings are in line with a few previous studies in Asia [18, 20–22], which also suggest that some Asian amphibian species and populations are more at risk of *Bd* infection than others [18]. Interestingly, the populations of the Odorous Frog (*O. graminea*) located in lower latitudes were free of *Bd*, while a population of this species in site#1 was detected to be infected (S1 Table). This suggests that infection patterns may not always have a taxon-specific context, and could be linked to other factors such as climate.

Our results suggest that site-specific environmental conditions are key drivers of *Bd* prevalence in amphibian adults. Our analysis demonstrated that mean temperature of warmest quarter and elevation are the most important factors influencing *Bd* infection in susceptible individuals (S4 Table). In sub-tropical regions, warm temperatures during the breeding season likely exceed the optimal range for *Bd* [69]. This could reduce pathogen survival and transmission among hosts [53, 70–72]. We also found that elevation is related to variations in temperature, and at higher elevation sites (>700 m), cooler temperatures during the warmest months may be a mitigating factor. In addition, *Bd* prevalence appears to be associated with the habitats of adult hosts, with multiple habitats resulting in higher infection risk [52, 73, 74]. Taken together, these findings suggest that environmental conditions at a given location may be one of the most important factors driving *Bd* prevalence in amphibians.

Our study included 76 opportunistically collected tadpoles of several endemic species, representing only a small sample size. Some authors indicate that most *Bd*-related studies have focused on post-metamorphic Asian frogs [26], suggesting that the role of metamorphic stages in the dispersal and persistence of *Bd* may be currently under-estimated [37, 75]. Our findings supports this notion; we observed that some tadpoles of the common species *H. guentheri* were frequently infected with *Bd* whereas adults were not, suggesting that *Bd* surveys should investigate infection patterns across anuran life-history stages in endemic species [26] in order to better assess infection dynamics.

### ITS-based haplotype diversity and distribution

Our phylogenetic analysis showed that the new haplotypes uncovered belonged to the earliest emerging clade in the tree, *Bd*ASIA-1. This indicates that southern China may be important in the origins and spread of *Bd. Bd*-Asian lineages were present at all sites where *Bd* was found, with *Bd*ASIA-1 and *Bd*ASIA-3 lineages co-occurring at three sites (Fig. 4). Among the amphibians tested and the sites surveyed, haplotype GX03 (*Bd*ASIA-3) was the most prevalent, suggesting the widespread presence of *Bd*ASIA-3 in Southeast Asia [23]. Moreover, the few clinical signs of *Bd* infection observed in Asian amphibians were mainly attributed to this haplotype (*Bd*ASIA-3 lineage) [76].

The limited presence of the *Bd*GPL lineage in this study is different from findings of previous studies in China, Japan, and Brazil, which revealed its widespread distribution [18, 21, 24, 77, 78]. High haplotype richness of the *Bd*GPL has been identified from native frogs dwelling in forested areas in India [67, 79]. Given the low genetic diversity, we suspect that *Bd*GPL in the studied sites may have been recently introduced to the region through anthropogenic activity, as has been observed elsewhere, due to its easy accessibility. Another possibility is that competition from endemic basal *Bd* lineages restricts the spread of *Bd*GPL.

Interestingly, the haplotype GX05 (*Bd*ASIA-1) occurs more frequently when co-occurring with GX01 (*Bd*GPL) and GX03 (*Bd*ASIA-3). This haplotype was detected at all sites and most species or populations. Its relatively strong competitive ability within microhabitats due to local conditions could potentially explain this prevalence pattern [80], although further investigation is needed.

Infected individual amphibians typically carried only a single haplotype, with the exception of one endangered tree frog that had two haplotypes (GX05 and GX06 - both from the *Bd*ASIA-1 lineage). While this is consistent with early studies in Japan [18, 21], numerous studies have since shown co-infection to be frequent in both native and invasive species [22, 24, 67]. In China, prior work has demonstrated that amphibians infected with *Bd* often carry multiple haplotypes, belonging to the *Bd*GPL lineage [24]. Similarly, introduced Northern American bullfrogs (*R. catesbeiana*) and endemic species (*Pelophylax nigromaculatus, F. limnocharis, Rana chaochiaoensis, Hyla annectans*) in China harbor *Bd*GPL lineages with many haplotypes co-occurring in the region [24]. In South Korea, many amphibians also carry multiple haplotypes of the endemic lineages of *Bd* (*Bd*ASIA-1 and *Bd*ASIA-2/BRAZIL). Further studies on haplotype co-occurrence in Asia are warranted as this could potentially lead to the unearthing of highly virulent and new genotypes [81, 82].

### Sub-lethal effects of chytridiomycosis to Asian amphibians

This study, for the first time in Asia, observed clinical signs of chytridiomycosis on a urodelean salamander (*P. inexpectatus*) in a natural habitat. The individual was found in a stream under a waterfall in southern China [83] and belongs to the Least Concern (LC) species category [25]. Despite the LC status, it was listed as a vulnerable species at the national level [84]. Our observation adds to the growing evidence that *Bd* infection can cause sub-lethal effects on some Asian amphibians, including the Bombay Night Frog (*Nyctibatrachus humayuni*) in the Western Ghats of India [76], Hasselt’s Toad (*Leptobrachium hasseltii*) in Indonesia [85], and the Japanese tree frog (*Hyla japonica*) [86]. Our observation associated with previous documented impacts of *Bd* in natural species populations in Asia, highlight that *Bd* infection might cause sub-lethal effects on some native amphibians, particularly in forest populations at high elevations, where population numbers are low owning to greater habitat fidelity and lower fecundity [87].

In conclusion, our research found new haplotype diversity in *Bd*-Asian lineages infecting isolated populations of previously unknown native amphibian species. Early emerging haplotypes closely related to *Bd*ASIA-1 and *Bd*ASIA-3 lineages, together with the global infection haplotype (*Bd*GPL), were identified, indicating southern China as a *Bd* diversity hotspot. Our findings also suggest that *Bd* prevalence in amphibians is affected by temperature and elevation within natural forests, and that it may pose sub-lethal effects on some susceptible amphibian species. These results reinforce the knowledge base for understanding the *Bd* infection dynamics of amphibians in China, in Asia, and globally.

## Supporting information

S1 Table. *Bd* prevalence of different sites in adult and larva amphibians

S2 Table. The ITS1-5.8S-ITS2 sequences of *Bd* used in this study.

S3 Table. The prevalence of *Bd* in months across 17 sites.

S4 Table. First 15 candidate models for *Bd* presence/absence in susceptible amphibian adults.

## Acknowledgements

We thank the support from Department of Forestry of Guangxi Zhuang Autonomous Region, and Guangxi Shiwandashan Natural Nature Reserve Administration, Guangxi Damingshan Natural Nature Reserve Administration, Guangxi Dayaoshan Natural Nature Reserve Administration and Guangxi Huaping Natural Nature Reserve Administration; Liam D. Fitzpatrick for advice in PrepMan method of DNA extraction; Changming Bai consultation in early research. We also thank the following people for their assistance with field work: Chenghai Fu, Shuyi Luo; Mingzhong Tao; Yongwen Lin; Zhoulin Tan; Shipeng Zhou; Yongqiang Cao; Cheng Fang; Amrapali P. Rajput; Jianjun Ou; Shubao Wei; Jianchun Li; Hankun Ling.

## Author Contributions

**Conceptualization:** Dan Sun, Gajaba Ellepola, Jayampathi Herath, Kris Murray, Madhava Meegaskumbura

**Data curation:** Dan Sun, Madhava Meegaskumbura

**Formal analysis:** Dan Sun, Gajaba Ellepola, Kris Murray, Madhava Meegaskumbura

**Funding acquisition:** Gajaba Ellepola, Jayampathi Herath, Madhava Meegaskumbura

**Investigation:** Dan Sun, Gajaba Ellepola, Jayampathi Herath, Hong Liu, Yewei Liu, Madhava Meegaskumbura

**Methodology:** Dan Sun, Gajaba Ellepola, Kris Murray, Madhava Meegaskumbura

**Project administration:** Madhava Meegaskumbura

**Resources:** Hong Liu, Madhava Meegaskumbura

**Supervision:** Kris Murray, Madhava Meegaskumbura

**Visualization:** Dan Sun, Gajaba Ellepola, Madhava Meegaskumbura

**Writing - original draft:** Dan Sun, Gajaba Ellepola, Jayampathi Herath, Hong Liu, Yewei Liu, Kris Murray, Madhava Meegaskumbura

**Writing - review&editing:** Dan Sun, Gajaba Ellepola, Jayampathi Herath, Hong Liu, Yewei Liu, Kris Murray, Madhava Meegaskumbura

